# Slot-like capacity and resource-like coding in a neural model of multiple-item working memory

**DOI:** 10.1101/181354

**Authors:** Dominic Standage, Martin Paré

## Abstract

For the past decade, research on the storage limitations of working memory has been dominated by two fundamentally different hypotheses. On the one hand, the contents of working memory may be stored in a limited number of ‘slots’, each with a fixed resolution. On the other hand, any number of items may be stored, but with decreasing resolution. These two hypotheses have been invaluable in characterizing the computational structure of working memory, but neither provides a complete account of the available experimental data, nor speaks to the neural basis of the limitations it characterizes. To address these shortcomings, we simulated a multiple-item working memory task with a cortical network model, the cellular resolution of which allowed us to quantify the coding fidelity of memoranda as a function of memory load, as measured by the discriminability, regularity and reliability of simulated neural spiking. Our simulations account for a wealth of neural and behavioural data from human and non-human primate studies, and they demonstrate that feedback inhibition not only lowers capacity, but also lowers coding fidelity by all three measures. Because the strength of inhibition scales with the number of items stored by the network, increasing this number progressively lowers fidelity until capacity is reached. As such, the model provides a mechanistic explanation for experimental data showing that working memory precision decreases with increasing memory load before levelling off at capacity. Crucially, the model makes specific, testable predictions for neural activity on multiple-item working memory tasks.

## Introduction

Working memory refers to the retention of information for use in cognitive tasks, over intervals on the order of seconds. Visual working memory (WM) is a particularly active research field, largely because the high precision of the visual system affords fine-grained measurements that address the storage limitations of WM. These limitations are highly correlated with measures of intelligence and are currently the subject of intense research interest [see Luck and Vogel (2013)].

For several decades, research on storage limitations was dominated by the hypothesis that WM is supported by a small number of discrete ‘slots’ Accordingly, information is either stored with high precision in a slot or is simply not encoded if the number of items *n* exceeds the number of slots [see Cowan (2001)]. More recently, evidence has emerged for an alternative hypothesis, according to which a limited ‘resource’ *R* is allocated to *n* items, with no limit on *n*. Accordingly, the precision of WM representations decreases with increasing *n*, since less resource is available for the encoding of each item, *i.e.* precision tracks *R/n*. Thus, the nature of WM storage limitations is fundamentally different under the slot and resource hypotheses, attributing constraints to capacity and resolution respectively. It is increasingly clear, however, that neither is complete [see Luck and Vogel (2013); Ma, Husain, and Bays (2014)]. Generally, the slot hypothesis (Slot) is over-constrained with respect to resolution, since it can’t account for a gradual decrease in precision with increasing *n* (Bays, Catalao, & Husain, 2009). Equally, the resource hypothesis (Resource) is over-constrained with respect to capacity, since it can’t account for a plateau in imprecision with a critical number of items, where this number appears to correspond to capacity (Zhang & Luck, 2008). Consequently, several hybrid hypotheses have been presented, accounting for data that can’t be explained by Slot or Resource alone (Zhang & Luck, 2008; van den Berg, Shin, Chou, George, & Ma, 2012).

The above work has been invaluable in characterizing the storage limitations of WM, but does not speak to its neural basis. WM is widely believed to be supported by ‘attractor states’ in neocortex, emerging from recurrent excitation and feedback inhibition in local circuits. Under this framework, recurrent excitation sustains neural firing in the absence of driving stimuli (persistent activity), while feedback inhibition prevents this activity from running away [see Wang (2001)]. If *R* is instantiated by the cortical tissue mediating a task-relevant feature domain, *e.g.* spatial location, then feedback inhibition necessarily constrains capacity, since WM items will compete for representational space [see Franconeri, Alvarez, and Cavanagh (2013)]. If so, *R* cannot be infinitely divisible.Rather, it will be allocated to *n* ≤*K*items, with capacity *K* determined by the properties of feedback inhibition, *eg.* its strength and breadth. In other words, the simultaneous encoding of an arbitrary number of WM items is incompatible with feedback inhibition between stimulus-selective neural populations, a fundamental principle of neural information processing. The application of these principles to Resource leads to a strong hypothesis: a decrease in precision with increased memory load must be limited by capacity.

Here, we test this hypothesis with a biophysically-based model of a local circuit in posterior parietal cortex (PPC), a cortical area extensively correlated with WM (Gnadt & Andersen, 1988; Todd & Marois, 2004; Palva, Monto, Kulashekhar, & Palva, 2010; Christophel, Hebart, & Haynes, 2012; Salazar, Dotson, Bressler, & Gray, 2012). Previous studies have used similar models to offer mechanistic explanations for capacity (Edin et al., 2009), precision (Almeida, Barbosa, & Compte, 2015) and their relationship (Wei, Wang, & Wang, 2012; Roggeman, Klingberg, Feenstra, Compte, & Almeida, 2013), but these studies did not explain precision under the principles of Resource. Rather, they equated imprecision with the ‘drift’ of item-encoding neural populations in cortical tissue. As such, they make different predictions than our model (see the Discussion). According to Resource, imprecision reflects the signal-to-noise ratio (SNR) of neural representations. We extend this hypothesis from SNR to coding fidelity more generally, measuring the regularity and reliability of simulated spiking activity. In doing so, we demonstrate and explain the deterioration of coding fidelity with increasing *n* under established statistical measures, where this deterioration levels off at a critical *n*. Thus, we offer a novel explanation for resource-like coding and its relationship with capacity, unifying a large body of neural and behavioural data, and making specific predictions for experimental testing.

## Methods

Our local-circuit PPC model is a network of simulated pyramidal neurons and inhibitory interneurons, connected by AMPA, NMDA and GABA receptor conductance synapses (Figure 1A). Synaptic connectivity within and between classes of neuron was structured according to *in vitro* data, including structured and unstructured components of the connectivity to pyramidal neurons from interneurons (Figure 1B, Parameter Values in Methods). We refer to the former and latter components as local and broad inhibition respectively.

**Figure 1 (preceding page):**
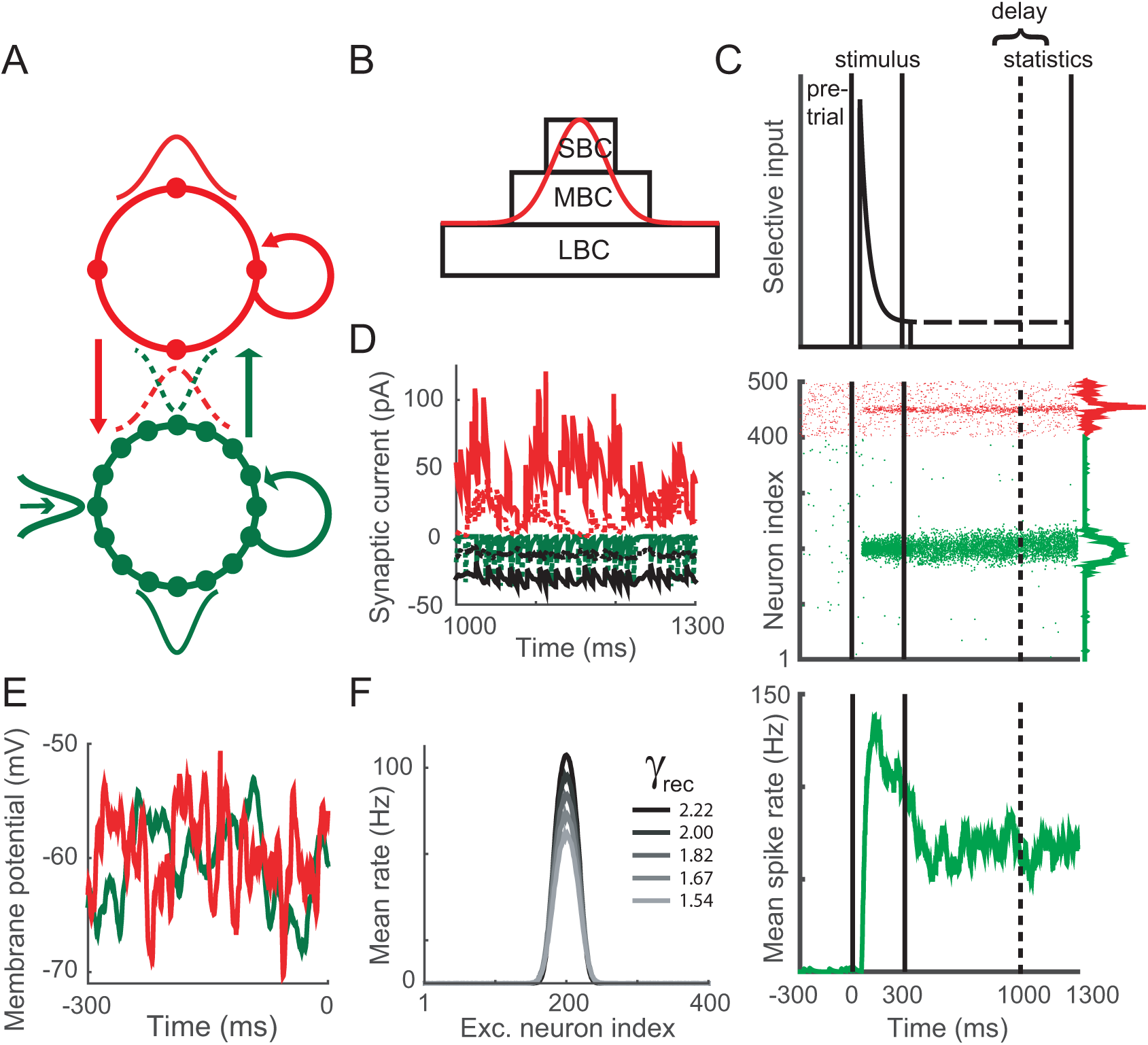
The local-circuit PPC model and simulated tasks. (A) Schematic of the model. Solid circles depict pyramidal neurons (green) and inhibitory interneurons (red), arranged periodically by their connectivity structures. The 4-to-1 ratio of pyramidal neurons to interneurons preserves their population sizes in the model. Arced and straight arrows depict synaptic connectivity within and between classes of neuron respectively. Thin Gaussian curves depict the structure of this connectivity (within, solid; between, dotted). The thick Gaussian curve depicts the RF of a pyramidal neuron. Red arrows depict GABAR synapses, the open green arrow depicts AMPAR-only synapses, and wide green arrows depict synapses with AMPARs and NMDARs. (B) Approximation of synaptic connections onto pyramidal neurons and interneurons from small, medium and large basket cells (SBC, MBC, and LBC respectively). Rectangles depict unstructured connectivity within each class of cell, and onto pyramidal neurons from each class. The red curve approximates their combined structure. (C) The visual and memory tasks are comprised of a pre-trial interval, stimulus interval and delay interval (top panel). Spiking statistics are taken during the last 300ms of the delay interval, referred to as the statistics window. Stimulus onset follows a 50ms visual response delay. On the visual (memory) task, stimuli persist (do not persist) throughout the delay interval, depicted by the dashed horizontal line. The decaying input signal simulates upstream response adaptation. An example trial of the 1-item memory task is shown in the middle and bottom panels. In the raster plot (middle panel), pyramidal neurons and interneurons are indexed from 1 - 400 and 401 - 500 respectively. Mean SDF (see text) over all pyramidal neurons and interneurons during the statistics window is shown to the right of the plot. Mean SDF over the item-encoding pyramidal population is shown in the bottom panel. (D) Synaptic currents onto a pyramidal neuron (solid) and an interneuron (dotted) during the delay interval of the 1-item memory task. Red, green and black curves show GABAR, AMPAR and NMDAR currents respectively. (E) Membrane potential of a pyramidal neuron and an interneuron during the pre-trial interval. (F) Mean rate over all pyramidal neurons during the statistics window of correct trials on the memory task for each value of control parameter *γ*_*rec*_ = 1/*γ*_*g*_ (see text).

We ran simulations of two common visual and WM tasks, a visually-guided delayed saccade task (the visual task) and a memory-guided delayed-saccade task (the memory task) [*e.g.* Paré and Wurtz (1997)]. Each task consists of three intervals: a pre-trial interval, a stimulus interval and a delay interval. Following the pretrial interval, items are presented during the stimulus interval on both tasks, remaining present during the delay interval on the visual task, but not the memory task (Figure 1C). We constrained the model by setting its parameter values according to anatomical and physiological data (Parameter Values in Methods), and by stipulating that it must qualitatively reproduce signature neural data from PPC (see Results, Figure 2). We then measured its storage capacity and coding fidelity as a function of *n*. Capacity was defined as the mean number of accurately encoded items during the last 300ms of the delay interval (the statistics window), where accurate encoding was determined by the rate, position (relative to stimulus position) and discriminability of item-encoding populations. We used three standard measures of coding fidelity: the SNR of stimulus-selective spiking, the coefficient of variation (CV) of interspike intervals (ISI), and the Fano factor (FF) of between-trial spike counts. SNR quantifies the degree to which selective spiking is discriminable from baseline activity, while CV and FF quantify the within-trial regularity and between-trial reliability of spiking respectively.

**Figure 2:**
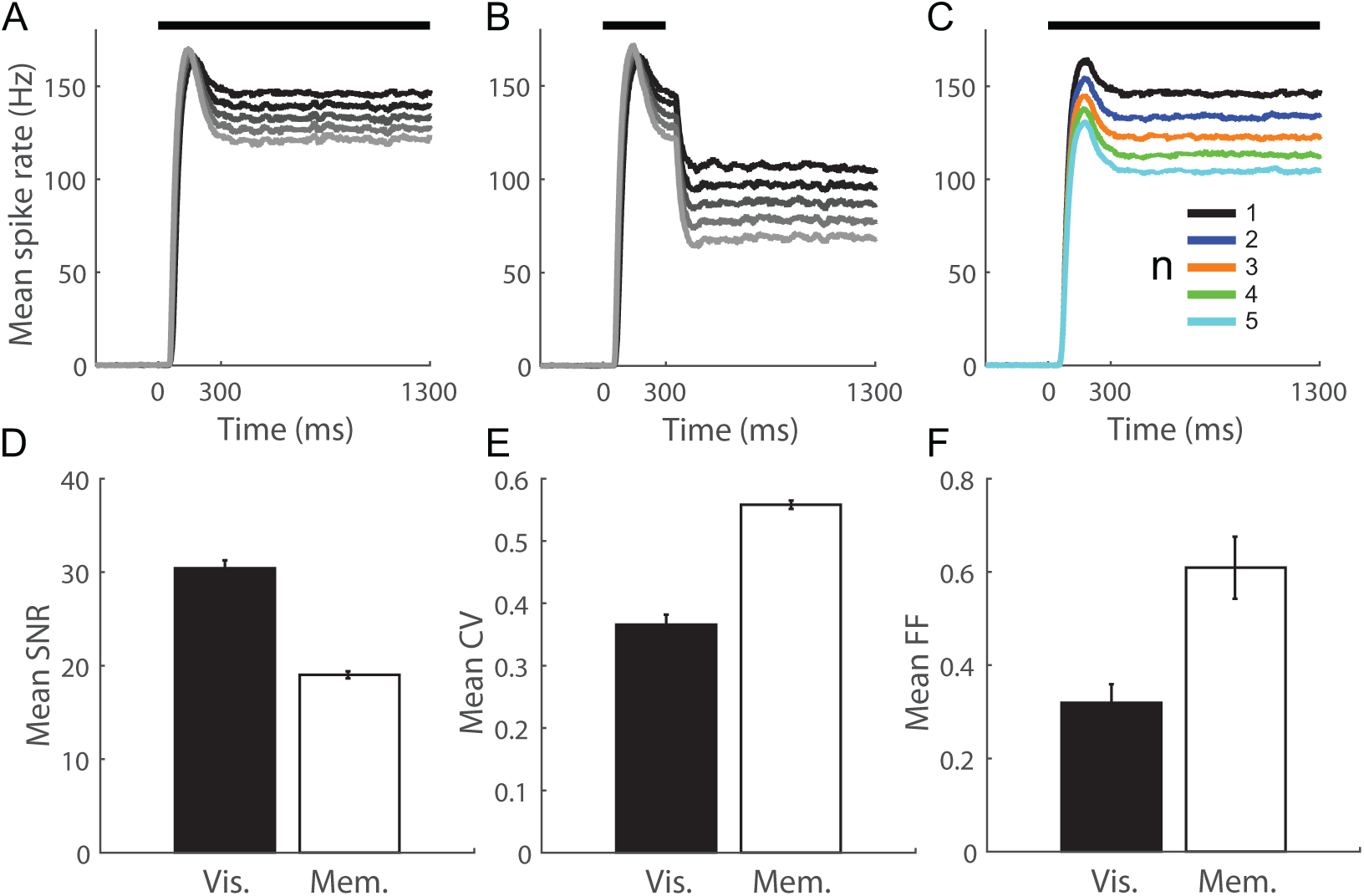
The model qualitatively reproduces signature neural data recorded from PPC during 1-item visual and memory tasks, and multiple-item visual tasks. (A) Mean activity at the RF centre of the item-encoding population on the 1-item visual task for each value of control parameter *γ*_*rec*_ = 1*/γ*_*g*_ (each gain condition, see text). Darker shades correspond to higher *γ*_*rec*_ (see legend in Figure 1F). (B) Mean activity at the RF centre on the 1-item memory task. (C) Mean activity at the RF centre of a single item-encoding population on the *n*-item visual task for all *n* (1 ≤ *n* ≤ 5). Results are shown for the highest gain condition. Thick horizontal bars show the timing of the target stimuli. (D-F) Persistent activity in the model encodes a low fidelity representation of an earlier stimulus, characterized by a lower SNR (D), higher CV (E) and higher FF (F) during the memory delay than the visual delay. Error bars show standard error of the mean. Results are shown for the lowest gain condition.

### The network model

The local circuit model is a fully connected network of leaky integrate-and-fire neurons (Tuckwell, 1988), comprised of *N*^*p*^ = 400 simulated pyramidal neurons and *N*^*i*^ = *N*^*p*^*/*4 fast-spiking inhibitory interneurons (putative basket cells). Each model neuron is described by

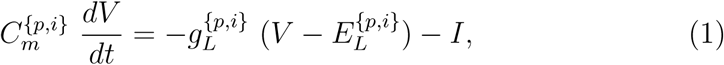

where *C*_*m*_ is the membrane capacitance of the neuron, *g*_*L*_ is the leakage conductance, *V* is the membrane potential, *E*_*L*_ is the equilibrium potential, and *I* is the total input current.When *V* reaches a threshold *ϑ*_*v*_, it is reset to *V*_*res*_, after which it is unresponsive to its input for an absolute refractory period of *τ*_*ref*_. Here and below, superscripts *p* and *i* refer to pyramidal neurons and interneurons respectively, indicating that parameter values are assigned separately to each class of neuron.

The total input current at each neuron is given by

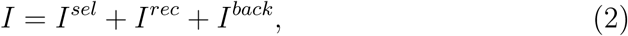

where *I*^*sel*^ is stimulus-selective synaptic current (set to 0 for interneurons), *I*^*rec*^ is recurrent (intrinsic) synaptic current and *I*^*back*^ is background current. *I*^*sel*^ and *I*^*rec*^ are comprised of synaptic currents, and *I*^*back*^ is comprised of synaptic current and injected current. Synaptic currents driven by pyramidal neuron spiking are mediated by simulated AMPA receptor (AMPAR) and/or NMDA receptor (NMDAR) conductances, and synaptic currents driven by interneuron spiking are mediated by simulated GABA receptor (GABAR) conductances. For AMPAR and GABAR currents, synaptic activation (the proportion of open channels) is defined by

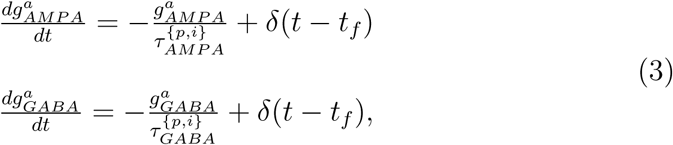

where *τ*_*AMP*__*A*_ and *τ*_*GABA*_ are the time constants of AMPAR and GABAR deactivation respectively, *δ* is the Dirac delta function, *t*_*f*_ is the time of firing of a pre-synaptic neuron and superscript *a* indicates that synapses are activated by different sources of spiking activity (selective, recurrent and background). NMDAR activation has a slower rise and decay and is described by

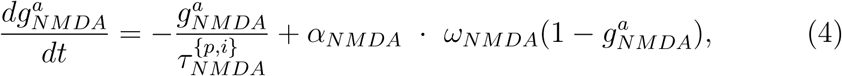

where *τ*_*NMDA*_ is the time constant of receptor deactivation and *α*_*NMDA*_ controls the saturation of NMDAR channels at high pre-synaptic spike frequencies. The slower opening of NMDAR channels is captured by

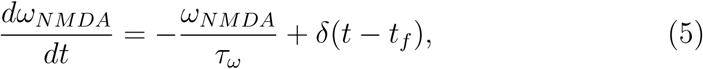

where *τ*_*ω*_ determines the rate of channel opening.

Intrinsic (recurrent, local feedback) synaptic current to each neuron *j* is defined by

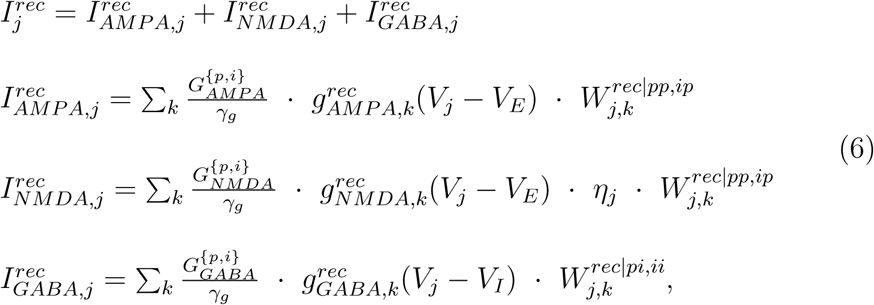

where *γ*_*g*_ is a scale factor controlling the relative strength of extrinsic and intrinsic synaptic conductance; *G*_*AMPA*_, *G*_*NMDA*_ and *G*_*GABA*_ are the respective strengths of AMPAR, NMDAR and GABAR conductance; *V*_*E*_ is the reversal potential for AMPARs and NMDARs, and *V*_*I*_ is the reversal potential *AM P A,k* for GABARs;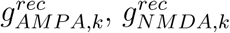, and 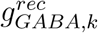 GABA,k are the activation of AMPAR, and *g* NMDAR and GABAR receptors respectively by pre-synaptic neurons *k*;*η* governs the voltage-dependence of NMDARs;and matrices *W* ^*rec*|*pp,ip*^ and *W* ^*rec*|*pi,ii*^ scale conductance strength or *weight* according to the connectivity structure of the network. This structure depends on the class of neuron receiving and projecting spiking activity, where superscripts *pp*, *ip*, *pi* and *ii* denote connections to pyramidal neurons from pyramidal neurons, to interneurons from pyramidal neurons, to pyramidal neurons from interneurons, and to interneurons from interneurons respectively. For each of these structures *s* ∈ {*pp, ip, pi, ii*}, *W* ^*rec*|*s*^ is a Gaussian function of the distance between periodically-arranged neurons, where the weight 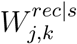 to neuron *j* from neuron *k* is given by

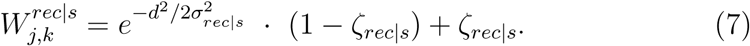

The distance between neurons is defined by *d* = *min*(❘*j*–*k* ❘ Δ*x* ^*p*^, 2 π – ❘*j* – *k*❘Δ*x*^*p*^) for *W*^*rec|pp*^, *d* = *min*(|*j* - *k*|_*x*^*i*^, 2π - |*j* - *k*|Δ*x*^*i*^) for *W*^*rec|ii*^, *d* = *min*(|*j*-*z*^*pi*^|Δ*x*^*p*^, 2π-|*j*-*z*^*pi*^|Δ*x*^*p*^) for *W* ^*rec*|*pi*^, and *d* = *min*(|*j*-*z*^*ip*^|Δ*x*^*i*^, 2π - |*j* - *z*^*ip*^|Δ*x*^*i*^)for *W* ^*rec*|*ip*^, with scale factors Δ*x*^*p*^ = 2*π/N*^*p*^ and Δ*x*^*i*^ = 2*π/N*^*i*^. For *W* ^*rec*|*pi*^ and *W* ^*rec*|*ip*^, *z*^*pi*^ = *N*^*p*^*/N*^*i*^ · *k* and *z*^*ip*^ = *N*^*i*^*/N*^*p*^*k* respectively. Parameter *σ*_*rec*|*s*_ determines the spatial extent of connectivity and parameter*ζ*_*rec*|*s*_ allows the inclusion of a baseline weight, with the function normalized to a maximum of 1 (0 ≤*ζ* rec|s*<* 1).

### Background activity

For each neuron, *in vivo* cortical background activity is simulated by current *I*^*back*^, defined by

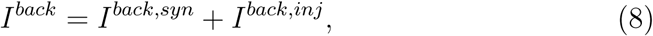

where *I*^*back, syn*^ is driven by synaptic bombardment and *I*^*back, inj*^ is noisy current injection. The former is generated by AMPAR synaptic activation, where independent, homogeneous Poisson spike trains are provided to all neurons at rate *µ*_*back*_. *I*^*back,syn*^ is therefore defined by

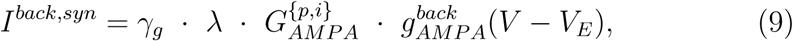

where *λ* is a scale factor and 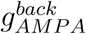 is given in Equation 3.

For *I*^*back,inj*^, we used the point-conductance model by (Destexhe, Rudolph, Fellous, & Sejnowski, 2001):

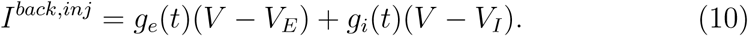

The time-dependent excitatory and inhibitory conductances *g*_*e*_(*t*) and *g*_*i*_(*t*) are updated at each timestep Δ*t* according to

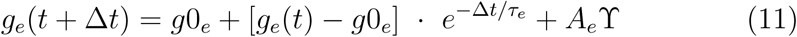

and

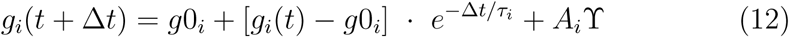

respectively, where 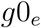 and 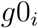 are average conductances, *τ*_*e*_ and *τ*_*i*_ are time constants, and γ is normally distributed random noise with 0 mean and unit standard deviation. Amplitude coefficients *A*_*e*_ and *A*_*i*_ are defined by

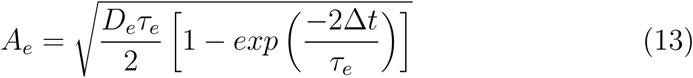

and

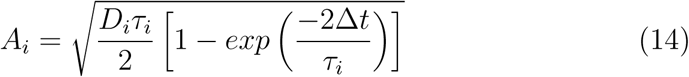

respectively, where 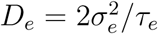 and 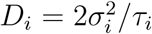 are noise ‘diffusion’ coefficients. See Destexhe et al. (2001) for the derivation of these equations.

## Experimental design and statistical analysis

### Simulated experimental tasks

We simulated the target stimuli in both tasks by providing independent, homogeneous Poisson spike trains to all pyramidal neurons *j* in the network, where spike rates were drawn from a normal distribution with mean *µ*_*sel*_ corresponding to the centre of a Gaussian response field (RF) defined by 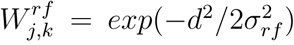. Constant d is given above for recurrent synaptic structure*W* ^*rec*|*pp*^, *σ*_*rf*_ determines the width of the RF and subscript *k* indexes the neuron at the RF centre. Spike response adaptation by upstream visually responsive neurons was modelled by a step-and-decay function

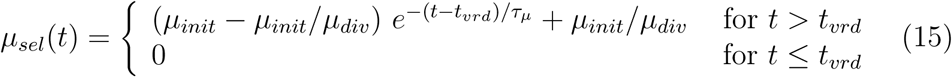

where *µ*_*init*_ determines the initial spike rate, *µ*_*div*_ determines the asymptotic rate, *τ*_*µ*_ determines the rate of upstream response adaptation, and *t*_*vrd*_ is a visual response delay. We simulated the visual task by providing these selective spike trains for 1300ms, following the 300ms pre-trial interval. We simulated the memory task by providing the selective spike trains for 300ms, following the pre-trial interval and followed by a 1000ms delay (Figure 1C). The stimuli were mediated by AMPARs only, so for all pyramidal neurons *j* in the PPC network,

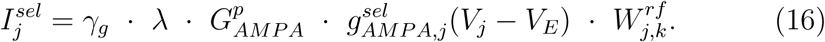

All simulations were run with the standard implementation of Euler’s forward method, where the time step was Δ*t* = 0.25ms.

### Determining working memory performance

We ran 400 trials of the visual and memory tasks with 1 - 5 stimuli (henceforth the *n*-item visual and memory tasks; 1 ≤ *n* ≤ 5). To determine WM performance on each trial of the memory task, spike density functions (SDFs) were calculated for all pyramidal neurons in the network by convolving their spike trains with a rise-and-decay function

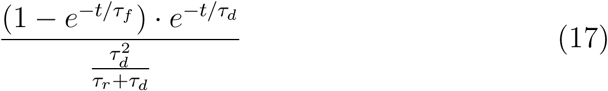

where *t* is the time following stimulus onset and *τ*_*r*_ = 1ms and *τ*_*d*_ = 20ms are the time constants of rise and decay respectively (Thompson, Hanes, Bichot, & Schall, 1996;Standage & Paré,2011).On each *n*-item trial, we calculated the mean of the SDFs over the last 300ms of the delay, obtaining the average activity over the network, and then partitioned the network into *n* equal regions. The location of each item was centred within each region. We then fit the mean activity in each region with a Gaussian function with four parameters: the height of the peak, the position of the peak, the standard deviation (controlling width), and the height that is approached asymptotically from the peak. An item was considered accurately stored if the fitted Gaussian satisfied three criteria: the height parameter exceeded 30Hz, the difference between the height and the fitted asymptote on both sides of the peak exceeded 15Hz, and the position parameter was within Δ*c* = 10 degrees of the centre of the RF for that item. For the first criterion, we chose 30Hz because this spike rate implies ∼ 10 spikes during the 300ms statistics window, as required to faithfully calculate CV and FF (Nawrot, 2010).The second criterion dictates that items are only considered accurately stored if the population response is discriminable. The third criterion ensures that the memory of the location of the item is close to the actual location, the precise value of which wasn’t crucial to our results (Δ*c >*∼ 5).

### Calculating spiking statistics

We selected *m* = 20 simulated pyramidal neurons from the network (the target neurons) and recorded their activity on *m* trials each. This population of neurons consisted of the neuron at the centre of the RF for a given target, and the *m* - 1 neurons closest to the RF centre. For each of the two tasks, the SNR of each target neuron was calculated on each trial by subtracting the spike count during the 300ms pre-trial interval from the spike count during the statistics window and dividing the result by the latter [*SNR* = (*SC*_*del*_ - *SC*_*pre*_)*/SC*_*pre*_, where *SC* is the spike count].

The CV of ISI was calculated for each target neuron on each trial by dividing the mean ISI by the standard deviation of ISI during the statistics window 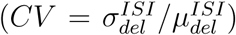The FF was calculated for each target neuron by recording the spike count during the statistics window on each trial, and dividing the variance by the mean over all trials for that neuron 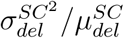These statistics were only calculated for accurately stored items, and from neurons that emitted at least 9 spikes during the 300ms statistics window (30Hz, see previous section). To increase statistical power on memory trials with *n >* 1 items, if the network did not accurately store the ‘first’ item, we searched for a corresponding neuron in another item-encoding population, where correspondence was determined relative to the RF centre, *e.g.* if the target neuron was located 3 indeces below the RF centre of item 1, we used a neuron located 3 indeces below the RF centre of another item. If no items were accurately stored, the trial was discarded for statistical purposes.

### Parameter values

In setting parameter values in the model, our aim was to justify every value by anatomical and physiological data, thus constraining our choices as much as possible, and then to use a single control parameter to explore the model’s performance and spiking statistics on the visual and memory tasks. Our control parameter was *γ*_*g*_, governing the relative strengths of extrinsic and intrinsic synaptic conductance and therefore the strength of recurrent processing.

For cellular parameters, we used standard values for integrate-and-fire neurons in cortical simulations (Compte, Brunel, Goldman-Rakic, & Wang, 2000), justified by electrophysiological data in earlier, related work (Troyer & Miller, 1997; Wang, 1999). These values are 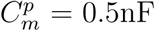, 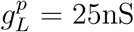, 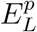 = −70mV, 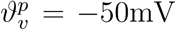, 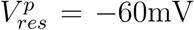 and 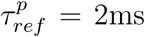; and 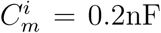, 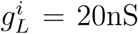, 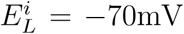, 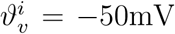, 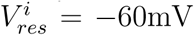 and 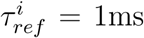. Likewise, synaptic reversal potentials are *V*_*E*_ = 0mV and *V*_*I*_ = -70mV, and the parameters governing the opening and saturation of NMDARs are *τ*_*ω*_ =2ms and *α*_*NMDA*_ = 0.5kHz respectively (Compte et al., 2000). The voltage-dependence of NMDARs is given by *η* = 1*/*[1 + *Mg · exp*(-0.062 · *V*)*/*3.57], where *Mg* = 1mM is the extracellular Magnesium concentration and *V* is measured in millivolts (Jahr & Stevens, 1990).

In setting parameters for the conductance strengths and time constants of decay of AMPARs and NMDARs, we followed Standage, You, Wang, and Dorris (2013), emphasising fast inhibitory recruitment in response to slower excitation [see Povysheva et al. (2006) for discussion]. For 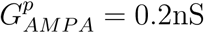, 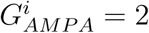. 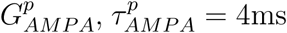 and 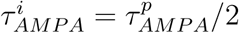; and for NMDARs, 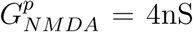, 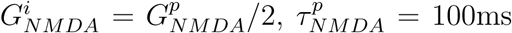 and 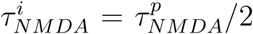. These values produce fast-decaying AMPAR currents on the order of 10pA (Angulo, Rossier, & Audinat, 1999; Desai, Cudmore, Nelson, & Turrigiano, 2002) that are stronger and shorter-lived onto inhibitory interneurons than onto pyramidal neurons (Hestrin, 1993; J. McBain & Fisahn, 2001; Hull, Isaacson, & Scanziani, 2009), and slow-decaying NMDAR currents on the order of 10pA (Berretta & Jones, 1996; Angulo et al., 1999) that are stronger and longer-lived at synapses onto pyramidal neurons than onto inhibitory interneurons (Hull et al., 2009).For 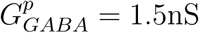and 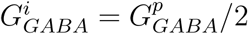 producing GABAR currents several times stronger than the above excitatory currents, where the stronger conductance at synapses onto pyramidal neurons captures their greater prevalence of GABARs (Markram et al., 2004). GABAR time constants were set 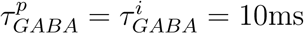 (Salin & Prince, 1996; Xiang, Huguenard, & Prince, 1998). Example synaptic currents are shown in Figure 1D.

The connectivity structures *W* ^*rec*|*pp,ip,pi,ii*^ capture the probability of lateral synaptic contact within and between classes of neurons in local cortical circuitry (Wilson & Cowan, 1973; Somers, Nelson, & Sur, 1995). A considerable volume of data indicates that the probability of lateral synaptic contact between cortical pyramidal neurons is normally distributed with mean0 and half-width of ∼ 0.25mm (Hellwig, 2000; Berger, Perin, Silberberg, & Markram, 2009; Voges, Schüz, Aertsen, & Rotter, 2010).Thus, *σ*_*rec*|*pp*_ corresponds to 0.25mm, determining the size of the cortical region being modelled, and *ζ*_*rec*|*pp*_ = 0.We are unaware of any data suggesting that the lateral projections of pyramidal neurons target basket cells differently than they target other pyramidal neurons, so we set *σ*_*rec*|*ip*_ = *σ*_*rec*|*pp*_ and *ζ*_*rec*|*ip*_ = *ζ*_*rec*|*pp*_. Arguably, *σ*_*rec*|*ip*_ should be narrower than *σ*_*rec*|*pp*_, since the dendritic trees of basket cells are less extensive than those of pyramidal neurons, but setting these parameters to equal values supported more stable network dynamics, *i.e.* it furnished sufficient local-circuit inhibition for the model to simulate the experimental tasks without modifications to other parameter values.

For connectivity structures *W* ^*rec*|*pi,ii*^, values for *σ*_*rec*|*pi,ii*_ and *ζ*_*rec*|*pi,ii*_ are justified by four premises: firstly, we assume that basket cells are a major source of lateral inhibition (Krimer & Goldman-Rakic, 2001) and we limit our focus to this class of inhibitory interneuron; secondly, basket cells synapse onto the somatic and perisomatic regions of their targets [see Markram et al. (2004)]; thirdly, the axons of basket cells contact their targets indiscriminately throughout the range of their ramifications (Packer & Yuste, 2011); and fourthly, the basket cell population can be divided into small (local arbour), medium (medium arbour) and large (wide arbour) cells in equal proportion, *i.e.* one third each (Krimer et al., 2005). Under the first and second premises, we do not need to consider the dendritic morphology of the targets of inhibitory interneurons, so we set *σ*_*rec*|*pi*_ = *σ*_*rec*|*pp*_. Under the second and third premises, we assume a uniform synaptic distribution for inhibitory targets, where the axonal ramifications of small, medium and large basket cells cover progressively larger areas (Krimer & Goldman-Rakic, 2001; Krimer et al., 2005), with large basket cells (LBC) covering the entire local circuit (Kisvá rday, Beaulieu, & Eysel, 1993; Markram et al., 2004). We therefore approximate this connectivity structure by setting *σ*_*rec*|*pi*_ = *σ*_*rec*|*ii*_ = 2.*σ*_*rec*|*pp*_ and *ζ*_*rec*|*pi*_ = *ζ*_*rec*|*ii*_ = 1*/*3, where the former corresponds to a half-width of∼ 0.5mm [*cf.* Kisvárday et al. (1993); Krimer and Goldman-Rakic (2001); Krimer et al. (2005)] and the latter refers to the 1*/*3 proportion of LBCs. This approach to determining inhibitory connectivity parameters is depicted in Figure 1B. We set *σ*_*rec*|*pp*_ = 0.2 because this value supported the simultaneous representation of 5 simulated visual stimuli, corresponding to the upper limit on human WM capacity, *i.e.* 4 ± 1 items (Luck & Vogel, 1997; Cowan, 2001). Finally, we set *σ* _*rec*|*ii*_ = *σ*_*rec*|*pi*_ because LBCs make extensive contacts onto one another over the full range of their axonal ramifications (Kisvárdayet al., 1993). Note that we do not attribute biological significance to the spatial periodicity of the network. Rather, this arrangement allows the implementation of *W* ^*rec*|*pp,ip,pi,ii*^ with all-to-all connectivity without biases due to asymmetric lateral interactions between neurons, and further captures the topographic mapping of spatially periodic stimuli in many visual [*e.g.* Thomas and Paré (2007)] and WM [*e.g.* Funahashi, Bruce, and Goldman-Rakic (1989); Matsuyoshi, Osaka, and Osaka (2014)] tasks. In Results section ’Feedback inhibition underlies slot-like capacity and resource-like coding’, we eliminated broad inhibition by setting *ζ*_*rec*|*pi*_ and *ζ*_*rec*|*ii*_ to 0, and we increased the strength of local feedback inhibition by setting 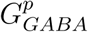 and *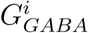* to 3.5nS and 1.75nS respectively.

In setting parameter values for background activity in each network, we initially omitted background synaptic input *I*^*back,syn*^ and followed the data by Fellous, Rudolph, Destexhe, and Sejnowski (2003) to produce *I*^*back,inj*^, where *g*0_*e*_ = 5nS and *g*0_*i*_ = 25nS, *τ*_*e*_ = 2.5ms, *τ*_*i*_ = 10ms, *σ*_*e*_ = 5nS and *σ*_*i*_ = 12.5nS. Because the average inhibitory background conductance *g*0_*i*_ is five times the average excitatory background conductance *g*0_*e*_ [see Destexhe (2010)], our simulated pyramidal neurons did not respond adequately to selective stimuli under these parameter values. We therefore reduced the average conductances by a factor of two, setting *g*0_*e*_ = 2.5nS, retaining the ratio of inhibitory to excitatory conductance strength *g*0_*i*_ = 5 *g*0_*e*_ = 12.5nS, and simulating the ‘other half’ of upstream cortical background activity by providing independent, homogeneous Poisson spike trains to all neurons in the network. As such, we assumed that each neuron forms ∼ 10, 000 synapses with upstream cortical neurons (Douglas, Markram, & Martin, 2004), and that by dividing *g*0_*e*_ and *g*0_*i*_ by two, we were effectively omitting ∼ 5, 000 background inputs. We therefore approximated 5000 upstream cortical neurons firing at 1Hz each by setting the rate of background Poisson spike trains to *µ*_*back*_ = 500Hz and setting the extrinsic synaptic scale factor to *λ* = 10, trading temporal summation for spatial summation (Prescott & De Koninck, 2003; Standage et al., 2013). As noted above, background spike trains were provided to all pyramidal neurons and interneurons in each network, mediated by AMPARs on the assumption that spike trains converging on PPC (an association cortical area) are predominantly ascending. Evidence for AMPAR-mediated ascending activity is provided by Self, Kooijmans, Supèr, Lamme, and Roelfsema (2012). This approach simultaneously released the network model from the overly-strong background inhibitory currents and implemented an established, biologically plausible form of gain modulation [balanced background inputs (Chance, Abbott, & Reyes, 2002)], rendering the PPC network responsive to simulated visual stimuli. Note that our parameter values for background current injection (*g*0_*e*_, *g*0_*i*_, *τ*_*e*_, *τ*_*i*_, *σ*_*e*_ and *σ*_*i*_) were based on recordings from pyramidal neurons (Fellous et al., 2003), but since we are unaware of any data to guide these parameters for inhibitory interneurons, we assigned them the same values for all neurons. The effect of this background activity on the membrane potential of a pyramidal neuron and an interneuron is shown in Figure 1E.

For the target stimuli, the width of RFs was determined by *σ*_*rf*_ = *σ*_*pp*_*/*2. This narrow width captures the less-extensive dendritic branching in cortical (input) layer 4 compared to layers 2/3 and 5 (see above for justification of lateral connectivity in the model). The initial spike rate at the RF centre was *µ*_*init*_ = 10, 000*/γ*_*g*_Hz, which (for *γ*_*g*_ = 1) can be equated with *e.g.* 100 upstream, visually-responsive neurons firing at 100Hz each, given our use of homogeneous, independent Poisson spike trains. Note, however, that the synaptic scale factor *λ* = 10 probably renders this spike rate unrealistically high, since it implies *e.g.* 1000 upstream neurons firing at 100Hz. Nonetheless, the high initial spike rate ensured a rapid-onset, high-rate visual response in the network for all processing regimes furnished by control parameter *γ*_*g*_, as observed experimentally [*e.g.* Paré and Wurtz (1997); Thomas and Paré (2007); Churchland, Kiani, and Shadlen (2008)]. Upstream, visual response adaptation was simulated by *µ*_*div*_ = 10 and *τ*_*µ*_ = 50ms. The former is somewhat extreme, but allowed the rate of the initial population response in PPC to exceed the steady state response on the visual task for all values of *γ*_*g*_ [*e.g* Paré and Wurtz (1997); Churchlandetal. (2008)].Our use of *γ*_*g*_ as a denominator in determining *µ*_*init*_ (Equation 15) supported stronger selective inputs when the network had stronger recurrent processing (smaller *γ*_*g*_), allowing the rapid-onset, high-rate visual response described above. For larger *γ*_*g*_, the network more readily gives way to its inputs, so a weaker input is sufficient to elicit a similar response. The visual response delay was *t*_*vrd*_ = 50ms (Thomas & Paré 2007).

## Results

To systematically investigate network performance on the visual and memory tasks, we varied a single parameter *γ*_*g*_, scaling the relative strength of intrinsic (Equations 6) and extrinsic (Equations 9 and 16) synaptic conductance. We ran a block of trials for a range of values of this parameter (increments of 0.05), searching for values supporting a mean capacity of atleast 0.95 items on the *n*-item memory task for *n* ≤ 5, and for which all item-encoding populations on the 5-item task co-existed at the end of the stimulus interval (with excessively strong intrinsic synapses, feedback inhibition produced strong competition between populations, so that not all populations were extant at the onset of the memory delay). Thus, we interpolated between upper and lower bounds on the strength of recurrent drive that support performance of the task, finding that our criteria were satisfied by *γ*_*g*_ ∈ {0.45, 0.5, 0.55, 0.6, 0.65}. We confirmed that these values support a stable background state (no structured activity before stimulus onset) by running a single trial with no stimuli for 10s, and that they support performance of the visual task (more than 99% of items were accurately encoded during the statistics window for all *n* and gain conditions). Because lower values of *γ*_*g*_ produce stronger recurrent drive and higher neuronal gain (Figure 1F), it is convenient to define *γ*_*rec*_ = 1*/γ*_*g*_. We refer to the values of *γ*_*rec*_ (equivalently *γ*_*g*_) as the gain conditions of the network. A single trial of the 1-item memory task is shown in Figure 1C.

### The model complies with signature neural data from PPC

Electrophysiological recordings from PPC show that on 1-item visual and memory tasks, the rate of stimulus-selective activity is higher during the visual delay than the memory delay; and on the memory task, the rate is higher during the stimulus interval than the memory interval (Paré & Wurtz, 1997). More generally, PPC activity consistently shows several characteristics across visual tasks, including a rapid-onset, high-rate response that drops to a steady state prior to movement-related activity [*e.g.* Paré and Wurtz (1997); Churchland et al. (2008); Louie, Grattan, and Glimcher (2011)], and a decrease in rate with an increase in the number of stimuli [*e.g.* Thomas and Paré (2007);Churchland et al. (2008); Louie et al. (2011)]. Consistent with these data, the mean rate of stimulus-selective spiking in the model was higher during the visual delay than the memory delay on the 1-item tasks (Figures 2A and B), and was higher during the stimulus interval than the delay interval on the memory task (2B). On the multiple-item visual tasks (2 ≤ *n* ≤ 5), selective spike rates were higher during the stimulus interval than the delay interval, and the rate of stimulus-selective activity decreased with increasing *n* (Figure 2C). These results were the case for all gain conditions, indicating that the model captured the relevant aspects of PPC processing over its full dynamic range.

For all gain conditions on the 1-item tasks, delay activity in the model had a lower SNR, higher CV and higher FF during the memory delay than the visual delay (Figures 2D, E and F). Thus, persistent activity encoded a low-fidelity representation of the stimulus, as reported in monkey PPC (Johnston et al, SfN abstracts, 2009). Higher gain conditions supported higher-fidelity encoding of the stimulus (see results for *n* = 1 in Figure 5C, D and E). Quantitative consideration of these results is provided in the Discussion.

**Figure 5 (preceding page):**
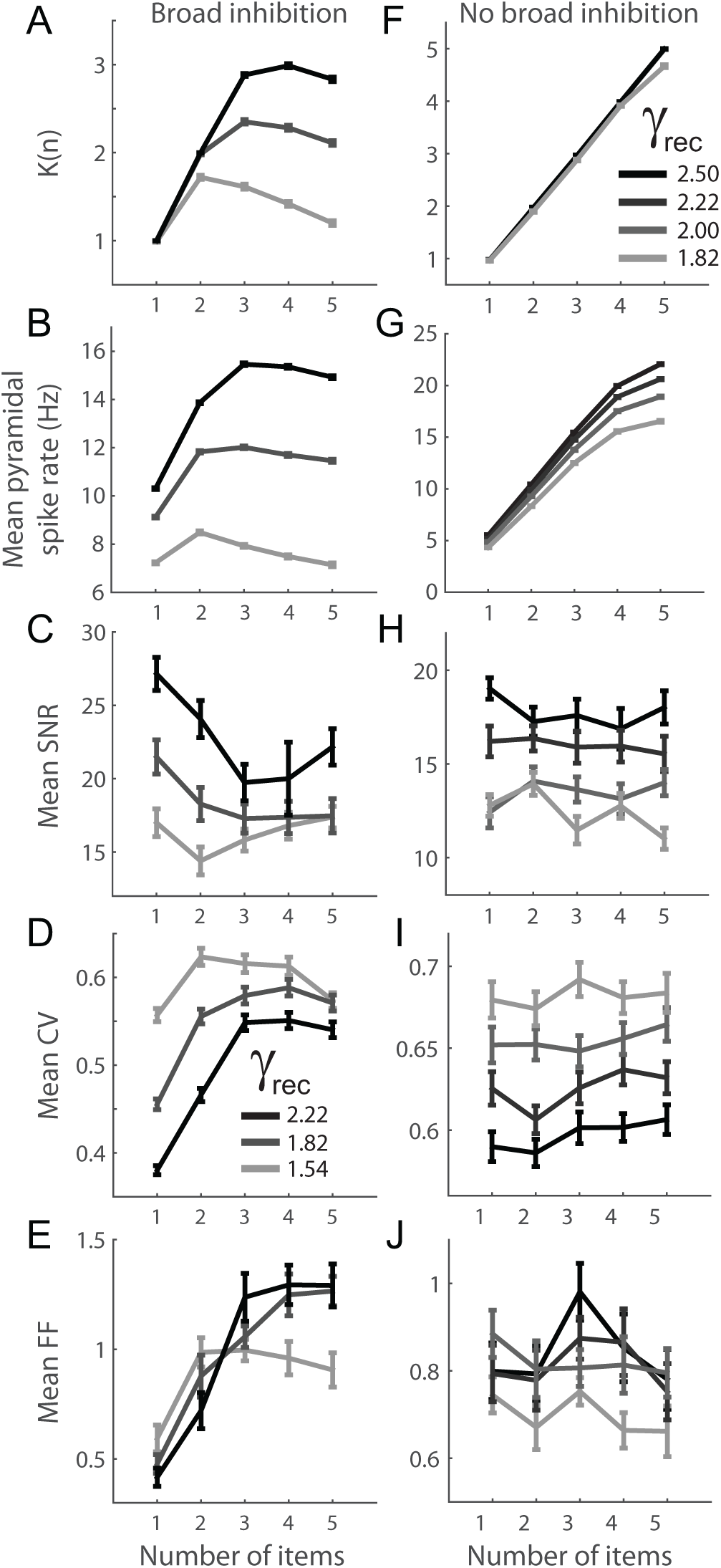
(A) The mean number of items accurately stored on each *n*-item memory task, referred to as capacity *K*(*n*). Error bars show standard error. Results are shown for the highest, median and lowest gain conditions, where darker shades correspond to higher gain (legend in panel D). Other gain conditions are omitted for clarity.(B) Mean spike rate of all pyramidal neurons in the model as a function of *n*. (C) SNR over *n*.(D) CV over *n*. (E) FF over *n*. (F-J) Removing broad inhibition increased capacity (*K*(*n*) ≈ *n* for all gain conditions, F), disrupted the correspondence between *K*(*n*) and the mean spike rate of all pyramidal neurons (G), and rendered SNR (H), CV (I) and FF (J) roughly independent of *n*. Darker shades correspond to higher gain conditions (legend in panel F), which were determined in the same way as in the original network (see text).

### Working memory performance in the model is consistent with that of monkeys and humans

To measure WM performance, we calculated the mean number of accurately stored items on each *n*-item memory task, referring to this quantity as capacity *K*(*n*). Example trials are shown in Figure 3.Mean spike rates at the RF centres of all accurate item-encoding populations are shown in Figure 4. Moderate to high gain conditions supported a maximum capacity (max[*K*(*n*)]) of around 2 and 3 items respectively (Figure 5A), consistent with WM capacity in monkeys (Heyselaar, Johnston, & Paré, 2011) and humans (Vogel & Awh, 2008; Luck & Vogel, 2013). In keeping with earlier models of this class (Edin et al.,2009;Wei et al., 2012), *K*(*n*) conformed to WM ‘overload’, decreasing beyond a critical *n* for all gain conditions (Matsuyoshi et al., 2014; Fukuda, Woodman, & Vogel, 2015). Finally, capacity was roughly tracked by the total pyramidal neuron activity in the network (Figure 5B), similar to electroencephalogram (EEG) recordings from PPC (Vogel & Machizawa, 2004).

**Figure 3:**
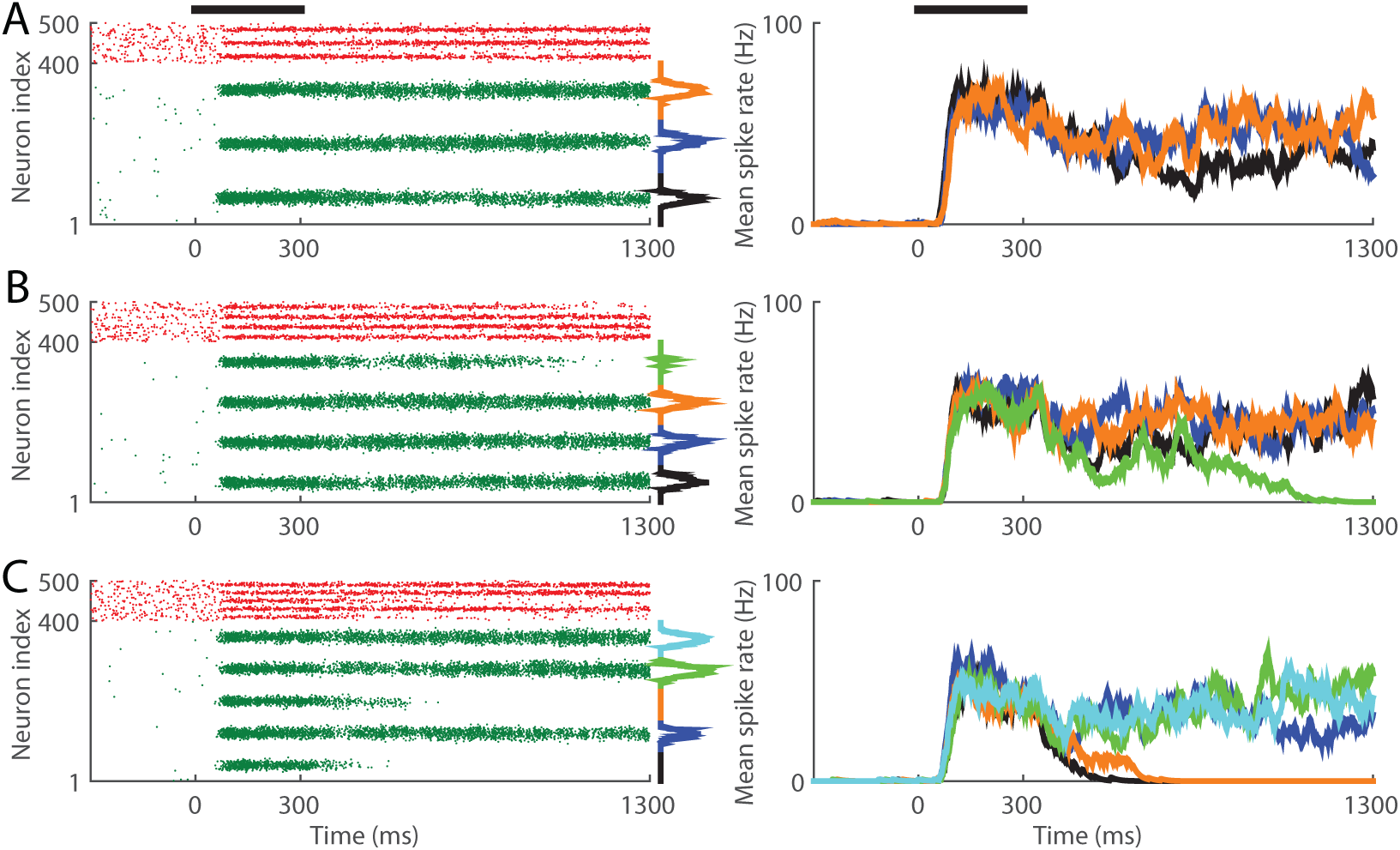
Three trials of the multiple-item memory task for the highest gain condition, with three (A), four (B) and five (C) items. The model accurately stored three items on each of these trials. Mean rates over all pyramidal neurons during the statistics window are inset to the right of the raster plots (see Figure 1C caption), where shades match the mean SDFs in the right-side panels. Thick horizontal bars in the top row show the timing of the stimuli.

### Predictions for experimental testing and their implications for Slot and Resource

Having demonstrated that our simulations are qualitatively consistent with a range of electrophysiological and behavioural data from visual and WM tasks, we investigated the model’s predictions for coding fidelity on multiple-item WM tasks, and the implications of these predictions for Slot and Resource. For all gain conditions, the coding fidelity of persistent activity on the memory task deteriorated as the number of items increased from *n* = 1 to *n* = 2, characterized by a decrease in SNR and an increase in CV and FF (Figure 5C-E). With higher gain (0.45 ≤ *γ*_*g*_ ≤ 0.55), for which *K*(3) *> K*(2), this reduction in coding fidelity continued as the number of items increased from *n* = 2 to *n* = 3, as measured by all three statistics. Within this range of *γ*_*g*_, coding fidelity leveled off as the number of items increased beyond *n* = 3, roughly tracking *K*(*n*) (Figure 5A). This finding is strikingly consistent with behavioural data showing that WM precision decreases with increasing *n* until it reaches a plateau at around 3 or 4 items (Zhang & Luck, 2008). It also suggests that the same mechanism is responsible for constraints on capacity and resolution: the competition between item-encoding populations, mediated by broad inhibition.

### Broad feedback inhibition underlies slot-like capacity and resource-like coding

If broad inhibition is responsible for slot-like capacity and resource-like deterioration of coding fidelity, then removing it will increase *K*(*n*) and eliminate the dependence of coding fidelity on *n*. We therefore removed the unstructured component of feedback inhibition, maintaining network stability by increasing the strength of local (structured) feedback inhibition (Parameter Values in Methods). We then determined the corresponding upper and lower values of *γ*_*g*_ according to the criteria above (top of Results),and repeated our simulations of the visual and memory tasks under these modified parameter values. As expected, these changes lead to a dramatic increase in *K*(*n*) (max[*K*(*n*)]*>* 4.6 for all *γ*_*g*_) (Figure 5F) and rendered coding fidelity roughly independent of *n* (Figure 5H-J).

The mechanism underlying resource-like coding is that a larger number of active item-encoding populations drives more broad inhibition, which reduces the (absolute) mean net current onto item-encoding neurons. This reduction in current decreases SNR for the simple reason that it decreases stimulus-selective spike rates (Figure 4), but pre-trial rates are fixed (constant of average). Indeed, this finding would be the case with earlier biophysical models of WM capacity (Edin et al., 2009; Wei et al., 2012) and precision (Wei et al., 2012; Roggeman et al., 2013; Almeida et al., 2015), since these models included broad inhibition (unstructured, uniform feedback inhibition). However, the reduction in current increases CV because the standard deviation of net current increases *relative to the mean* (Figure 6A). In other words, CV (the coefficient of variation of inter-spike intervals) reflects the coefficient of variation of net current (Pearson correlation coefficient *r >* 0.93 for all gain conditions). This increase in the relative variability of net current within each trial entails an increase in the relative variability of total net current across trials (*r >* 0.88 for all gain conditions), manifest as an increase in the relative variability of spike counts and therefore FF. In other words, FF (the Fano factor of spike counts) reflects the Fano factor of across trial total net current (Figure 6B). Of course, this explanation of FF assumes a tight correspondence between the total net current during the statistics window and the spike count, which is indeed the case (*r >* 0.97 for all gain conditions).

**Figure 6 (preceding page):**
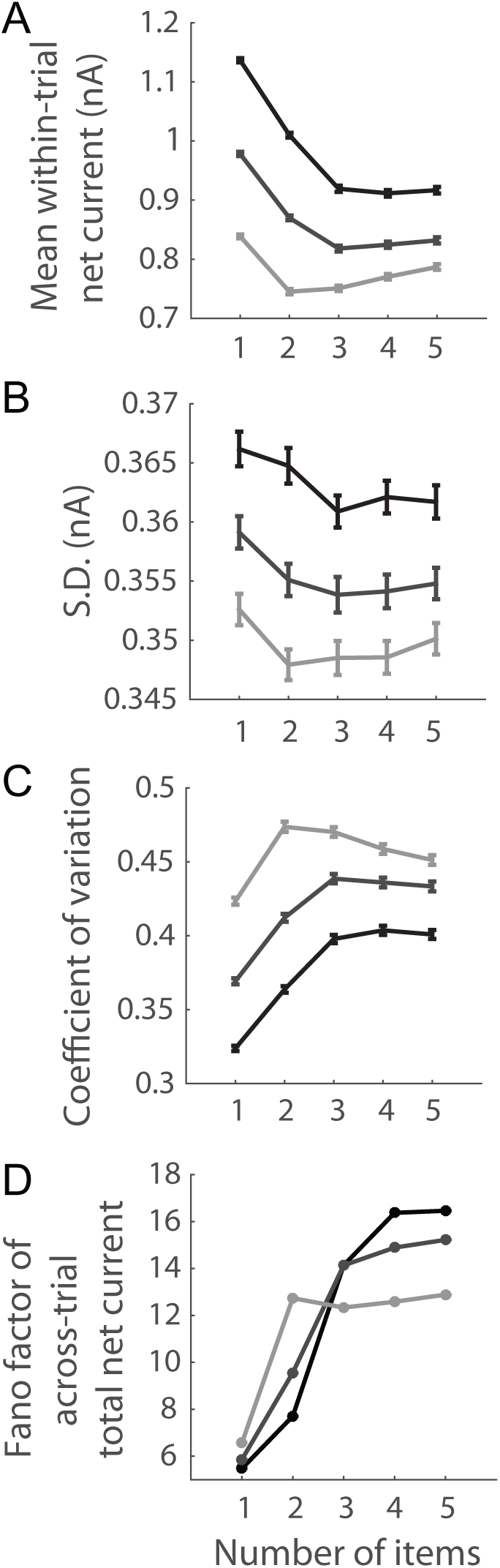
Inhibition raises CV and FF by increasing the relative variability of net current. Within each trial, the standard deviation of net current increases relative to the supra-threshold mean (A, rheobase=0.5nA for all *n* and gain conditions), so the higher coefficient of variation of net current de-regularizes spike timing to a greater degree. This increase in the variability of within-trial net current entails an increase in the variability of total net current on each trial (Pearson’s *r >* 0.88 for all gain conditions), resulting in an increase in the Fano factor (variance divided by the mean) of across trial total net current (B). Given the monotonic relationship between the total net current and the spike count (*r >* 0.97 for all gain conditions), the FF (of spike counts) reflects the Fano factor of total net current. (A) Mean (top), standard deviation (middle) and coefficient of variation (bottom) of within-trial net current at the RF centre of an item-encoding population during the statistics window (highest, median and lowest gain conditions). (B) Fano factor (variance divided by the mean) of the acrosstrial total net current for each *n* (highest, median and lowest gain conditions).

**Figure 4:**
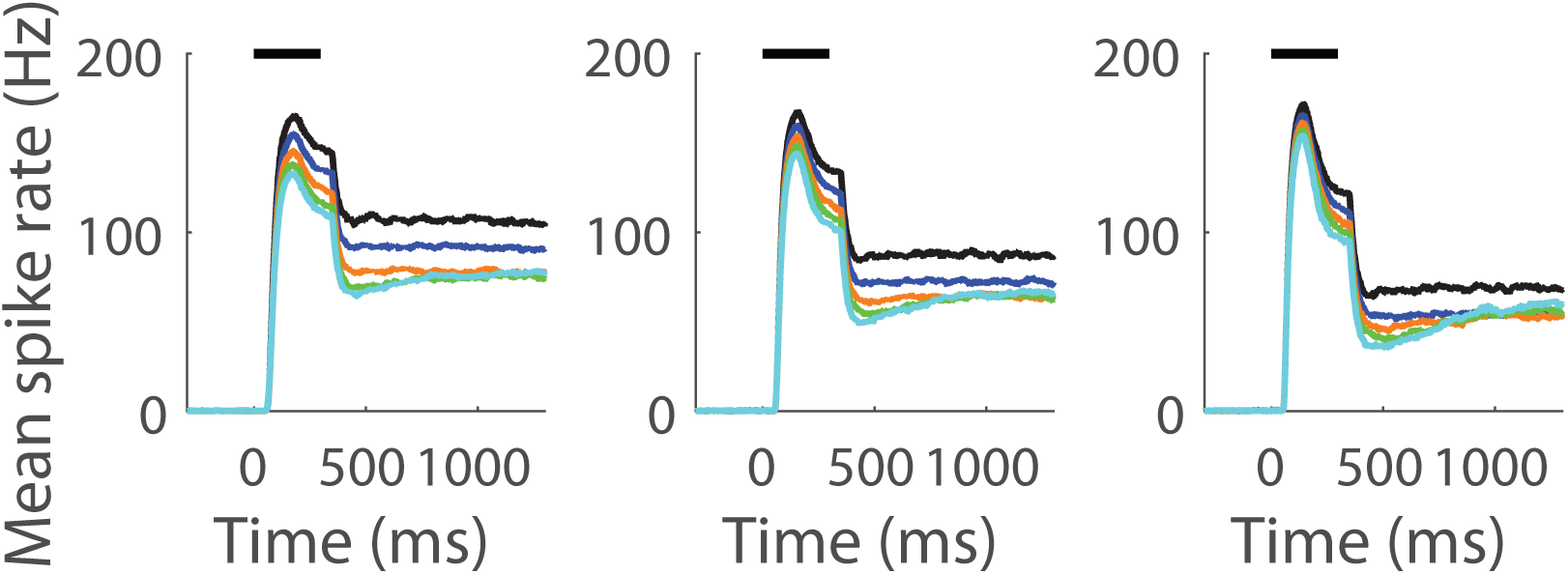
Mean rates at the RF centres of all accurate item-encoding populations on the memory tasks for each *n* (see legend in Figure 2C). Results are shown for the highest (left), median (middle) and lowest (right) gain conditions. Horizontal bars show the timing of the target stimuli.

## Discussion

Our local-circuit PPC model provides an integrated, mechanistic explanation for slot-like capacity and resource-like coding. Both are consequences of broad inhibition, which limits capacity by imposing competition (Edin et al., 2009), and reduces coding fidelity by lowering spike rates and rendering neurons more sensitive to current fluctuations. The model makes testable predictions for electrophysiological studies of WM. Most prominently, it predicts that on multiple-item tasks, the SNR (CV and FF) of persistent activity will decrease (increase) with increasing *n* until capacity is reached, levelling off thereafter. These predictions (Figure 5C-E) are strikingly consistent with behavioural data showing that WM precision decreases with increasing *n* until capacity is reached, plateauing thereafter (Zhang & Luck, 2008) (bilinear precision). They also explicitly demonstrate the incompatibility between Resource and mutual inhibition. The latter dictates that coding fidelity cannot decrease indefinitely with increasing memory load. Rather, it must be limited by competition. In this regard, we do not suggest that competitive dynamics in WM circuitry are immutable, dictating rigid capacity limitations. Far from it, we consider context-dependent control of neural dynamics to be fundamental to cognition [see Standage, Blohm, and Dorris (2014)], a view supported by recent studies in relation to WM storage (Edin et al., 2009; Roggeman et al., 2013; Almeida et al., 2015). Thus, we expect that capacity and precision will fluctuate with task-demands, but that their inherent relationship will hold: imprecision must be limited by capacity. Our findings offer a neural mechanism for this relationship.

### Quantitative considerations of coding fidelity

We have focused on the qualitative effect of memory load on coding fidelity, *i.e.* the direction of change in SNR, CV and FF as a function of *n*, but quantitative considerations warrant further comment. In particular, coding fidelity in our model was somewhat high with low *n* according to all three measures (Figure 5C-D). SNR is explained by low pre-trial rates (mean *<* 1Hz for all *γ*_*g*_), due to the high ratio of inhibitory to excitatory conductance in our method of background current injection. Our parameter values were determined by *in vivo* cortical data (see Methods section Parameter Values) and the low pre-trial rates they engender are consistent with neurons in the output layers of monkey primary visual cortex, in which spontaneous activity has been thoroughly investigated (Snodderly & Gur, 1995; Gur, Kagan, & Snodderly, 2005; Gur & Snodderly, 2008) [see also Maier, Adams, Aura, and Leopold (2010)]. In rodents, spontaneous activity is known to depend on intrinsic neuronal properties and connectivity, and differs between cortical layers [see Harris and Mrsic-Flogel (2013)]. Unfortunately, there is a lack of such data from extra-striate and association areas, an issue that should be addressed by future neurophysiogical studies. Our focus on coding fidelity concerns task-related spiking and we do not further pursue background activity here. Suffice to say, higher background rates would lower SNR in the model. As for CV and FF, it has long been maintained that their values should be around 1 *in vivo*, per the assumption that cortical spiking is Poissonian [see *e.g.* Shadlen and Newsome (1998)]. More recent data and analyses cast doubt on this assumption, as these measures are sensitive to the finite time windows from which they are calculated, non-stationary spike rates, and serial correlations in spike timing [see Nawrot, Boucsein, Molina, Aertsen, and Rotter (2008); Farkhooi, Strube-Bloss, and Nawrot (2009); Nawrot (2010); Rajdl and Lansky (2014)]. Of particular note, failure to account for within-trial fluctuations in spike rate (*e.g.* due to sensory stimuli) can lead to over-estimates of CV (Maimon & Assad, 2009;Nawrot, 2010), and between-trial fluctuations in rate (*e.g.* due to attentional state) or an insufficient number of spikes (less than ∼ 5 - 10) can lead to over-estimates of FF (Nawrot et al., 2008). It is worth noting that CV has been reported to be as low as ∼ 0.5 in PPC (Maimon & Assad, 2009), while FF has been reported in the range of ∼ 0.3 - 0.4 in visual cortex (Gur, Beylin, & Snodderly, 1997;Kara, Reinagel, & Reid, 2000).These are important issues for the understanding of non-task-related activity and neural coding, but none of them impacts our explanation of WM capacity and precision, nor its qualitative predictions.

WM capacity in our model also warrants further comment. As noted above, max [*K*(*n*)] was 2 or 3 items in moderate to high gain conditions (Figure 5A, consistent with data from non-human primate (NHP) and human subjects (Heyselaar et al., 2011; Luck & Vogel, 2013). Human studies have reported WM capacity higher than 3 items though [see Cowan (2010)]. Increasing the strength and decreasing the width of recurrent excitation readily increases capacity in our model, but these modifications do not change the finding that coding fidelity tracks capacity by all three measures used (not shown). Thus, the specific value of *K*(*n*) in each gain condition is parameter-dependent, but our chosen parameter values are consistent with the majority of experimental data on capacity [see Luck and Vogel (2013)]. These values are justified in Methods section Parameter values.

### Behavioural data accounted for by Slot and Resource

To a great degree, the conclusions drawn about WM storage limitations from experimental data depend on the nature of WM tasks and the ways in which performance is measured. For our purposes, WM tasks can be divided into two classes, referred to below as categorical and continuous report tasks. In both classes, information provided in a stimulus array must be retained over a delay interval. On categorical report tasks, performance is measured according to an all-or-none report on that information, such as whether the value of a particular feature is unchanged in a subsequent, post-delay stimulus array [*e.g.* Luck and Vogel (1997)]. On continuous report tasks, subjects report the memory of an analogue feature value [*e.g.* Wilken and Ma (2004)].

Slot accounts for behavioural data showing bilinear capacity on categorical report tasks (Luck & Vogel, 1997), *i.e.* for capacity *K*, subjects retain *n* items for *n* ≤ *K* and they retain *K/n* items for *n > K*. Slot also accounts for EEG (Vogel & Machizawa, 2004) and functional magnetic resonance imaging (fMRI) (Linden et al., 2003; Todd & Marois, 2004) data showing bilinear signal amplitude, where these signals correlate with *K*(*n*) (see below). On continuous report tasks, Slot cannot account for data showing decreasing precision with increasing *n* without recourse to the resource framework. Recent work has therefore referred to ‘discrete resource’ and ‘continuous resource’ hypotheses [see Fukuda, Awh, and Vogel (2010)]. The underlying premise of the former is that slots are a kind of resource that are used in a quantized manner, *i.e.* discrete sub-units of slots can be allocated flexibly to memoranda [*e.g.* Zhang and Luck (2008)]. This hybrid approach accounts for bilinear precision, since according to this hypothesis, the sub-units would be spread more thinly with increasing *n*, but all would be occupied for *n > K*.

Conversely with Slot, Resource cannot account for bilinear capacity on categorical report tasks, but accounts for monotonically decreasing precision with increasing *n* on continuous report tasks (Bays et al., 2009). In its original form (described above), Resource cannot account for bilinear precision, but can do so with the addition of trial-to-trial noise in the amount of resource allocated to each item (van den Berg et al., 2012). Such trial-to-trial variability has long been employed by hypotheses on cognition [*e.g.* perceptual decision making (Carpenter & Williams, 1995; Brown & Heathcote, 2005)] and does not deviate in principle from the original Resource formulation. Thus, we do not consider this approach to be a hybrid one. However, the bilinear signal amplitude shown by EEG (Vogel & Machizawa, 2004) and fMRI (Todd & Marois, 2004) studies on categorical report tasks poses a fundamental challenge to the resource framework, which can only account for these data if the relevant resource can be continuously *and partially* allocated to memoranda [see Fukuda et al. (2010)].

### Neural models of WM storage limitations

Abstract models instantiating Slot and Resource have been invaluable in characterizing WM storage limitations (Zhang & Luck, 2008; van den Berg et al., 2012), but they do not speak to the neural mechanisms that may implement their principles. In particular, these models do not account for persistent activity, widely believed to instantiate WM storage [for discussion, see Curtis and Lee (2010); D’Esposito and Postle (2015); Riley and Constantinidis (2016); Christophel, Klink, Spitzer, Roelfsema, and Haynes (2015)]. A number of studies have used implementation-level models to address the neural basis of capacity (Lisman & Idiart, 1995; Raffone & Wolters, 2001; Tanaka, 2002;Macoveanu, Klingberg, & Tegnér, 2006;Edin et al.,2009; Wei et al., 2012; Rolls, Dempere-Marco, & Deco, 2013). These models can be divided into two classes, attributing capacity to fundamentally different mechanisms. In one class, WM items are stored in oscillatory subcycles (*e.g.* beta-gamma oscillations nested inside alpha-theta oscillations), where different phases effectively isolate memoranda from one another (Lisman & Idiart,1995; Raffone & Wolters, 2001). As such, capacity is limited by the ratio of high-frequency to low-frequency oscillations. This compelling possibility relates feature-binding more broadly to WM (Raffone & Wolters, 2001), accounting for the finding that capacity does not depend on the complexity of WM items (Luck & Vogel, 1997;Awh, Barton, & Vogel, 2007) [though see Alvarez and Cavanagh (2004); Brady, Konkle, and Alvarez (2011)]. Notably, phase separation is fully consistent with the notion of discrete slots, and further accounts for data showing that different phases of gamma oscillations contain information about different memoranda (Siegel, Warden, & Miller, 2009). It is unclear, however, whether simultaneously-presented items could be allocated different phases. If not, these models imply that different WM mechanisms may encode simultaneously-presented and sequentially-presented memoranda.

In the other class of implementation-level model, multiple WM items are stored by attractor states (Tanaka, 2002; Macoveanu et al., 2006;Edin et al., 2009; Wei et al., 2012; Rolls et al., 2013; Roggeman et al., 2013; Almeida et al., 2015) *i.e.* the balance between recurrent excitation and feedback inhibition allows a limited number of memoranda to co-exist over a delay interval. A major difference between models of this class is the structure of local-circuit connectivity, where different connectivity structures embody different assumptions about the circuitry being simulated. In relation to capacity, the upshot of these studies is that feedback inhibition necessarily limits capacity [see Edin et al. (2009) for analysis], but the degree to which it does so can be ameliorated by mechanisms that localize and strengthen recurrent excitation.

Several of these studies also considered the neural basis of WM precision, equating imprecision with ‘drift’, or the deviation of item-encoding populations from the target locations (Wei et al., 2012; Roggeman et al., 2013; Almeida et al., 2015). In our model, drift actually decreased with *n* until capacity was reached, but its magnitude was negligible. The mean difference between drift on the 1-item task and on the *n*-item task corresponding to maximum capacity was Δ *d* = 0.38 degrees across gain conditions (maximum 0.52 degrees), calculated as the mean difference between the position parameter (see Methods) and the target location over all accurately encoded items, where ‘accuracy’ was re-calculated with *C* = 360*/n/*2. This re-calculation provided maximum tolerance for deviations, allowing up to 180 degrees with 1 item, 90 degrees with 2 items, and so on. Thus, our model predicts that drift has little bearing on WM precision under the task conditions simulated here. Interestingly, this prediction was also made by Almeida et al. (2015), whose model showed no significant drift as memory load was increased from 1 to 4 equidistant targets (the range considered by their model). Their behavioural data confirmed this prediction for 3 and 4 equidistant targets (the range considered by their experiments). Our model predicts that precision will decrease from 1 to 3 items, before levelling off at higher *n*, not because of drift, but because coding fidelity deteriorates until capacity is reached (3 items in this case). This discrepancy makes for good science: two models offer different mechanistic explanations for the same data, but make a different, testable prediction for a future experiment. Running this experiment would provide important evidence for one hypothesis over the other, but a definitive test requires neural data. Drift can be tested by constructing tuning curves from electrophysiological recordings of persistent activity on multiple-item WM tasks. To the best of our knowledge, no studies have done so, but we are aware of one study to quantify drift in this way from single-item WM data (Wimmer, Nykamp, Constantinidis, & Compte, 2014). These authors reported no appreciable drift in the average tuning bias of prefrontal cortical neurons prior to ∼ 2s (their Figure 3c). Thus, these data are in agreement with our findings over the timescale considered here. Note that our results were qualitatively unchanged with a 2s memory delay (not shown) and population width and drift remained severely limited (mean Δ*d* = 0.44 degrees). The above differences between neural models largely reflect their connectivity structures (our parameter values are justified in Methods). For example, inhibition was unstructured in the model by Wei et al. (2012) (broad inhibition only), so total pyramidal activity was invariant over memory load, a finding that is fully consistent with Resource. In our model (with structured and unstructured inhibition), total pyramidal activity during the statistics window tracked *K*(*n*) on the memory task (Figure 5A and B), as did the total (absolute) synaptic current (not shown). Our model is therefore consistent with bilinear EEG (Vogel & Machizawa, 2004) and fMRI (Todd & Marois, 2004) signals respectively. Ultimately, WM storage limitations may reflect constraints on the encoding, maintenance and/or decoding of memoranda (Ma et al., 2014). Our results emphasise the discriminability, regularity and reliability of persistent spiking as sources of WM imprecision, thereby implicating maintenance and decoding; but we do not suggest that these factors are the only sources of imprecision. Crucially, our study uses established measures of coding fidelity for single-neuron data (SNR, CV and FF), so our predictions for coding fidelity on multiple-item tasks are testable with single-neuron recordings.

### Limitations of the model

Of course, our model has limitations. To begin with, it only considers the spatial location of memoranda, ignoring other features and their conjunctions. In effect, our simulations assume that everything encoded by PPC satisfies a set of rules for selection. This approach is common among attractor models [*e.g.* Tanaka (2002); Macoveanu et al. (2006); Edin et al. (2009); Wei et al. (2012); Almeida et al. (2015)] and is justified in studies focused on storage limitations, *i.e.* it focuses on the mutual influence of persistently-active neural populations, regardless of the features or rules that lead to their initial activation. As noted above, models in which memoranda are stored in oscillatory subcycles can account for feature binding with sequentially presented stimuli (Lisman & Idiart, 1995; Raffone & Wolters, 2001). A more general understanding of feature-bound memoranda will likely require hierarchical models with converging feature maps. Such models have the further potential to explain the neural basis of ‘ swap errors’, where subjects report the feature value of a WM item other than the one probed [see Bays (2016)]. Hierarchical models also have the potential to explain the flexibility of WM precision, that is, the finding that one item can be maintained with higher precision than the others, but at a cost to those other items [see Ma et al. (2014)]. This finding points to the relationship between WM and attention, and to the modulation of item-encoding populations in distributed circuitry. These and related phenomena are beyond the scope of the present study, the focus of which is the mechanistic relationship between capacity and precision.

Finally, it is worth noting that our study is part of an ongoing research program investigating the neural basis of WM storage limitations with NHP subjects, in which *in vivo* electrophysiological recordings can be made during WM tasks. Thus, our simulations are purposefully constrained by our experiments, *e.g.* our use of equidistant targets [see Heyselaar et al. (2011) for justification]. This approach facilitates the testing of our predictions for single-neuron activity. In recent years, several studies have used categorical report tasks with NHP subjects [*e.g.* Heyselaar et al. (2011); Elmore et al. (2011); Lara and Wallis (2012)], but we are unaware of any studies to use continuous reports tasks with a non-human species. Future work must address this challenging gap.

## Conclusions

While recent studies have investigated neural mechanisms for WM capacity and precision (Wei et al., 2012; Roggeman et al., 2013; Okimura, Tanaka, Maeda, Kato, & Mimura, 2015), to the best of our knowledge, no previous study has proposed a neural mechanism for precision under the principles of Resource that also accounts for persistent activity Studies have proposed that Resource is implemented by the gain of item-encoding populations (van den Berg & Ma, 2014) and by divisively-normalized probabilistic spiking (Bays, 2014), but these studies have taken persistent activity for granted,*i.e.* they did not address the mechanisms by which an unlimited number of low-gain or divisively-normalized neural populations would be sustained over a memory delay. As described above, nested oscillations and attractor dynamics constrain capacity, so other mechanisms would be required. Our simulations without broad inhibition (Figure 5F-J) are instructive in this regard, since limiting competition rendered coding fidelity independent of memory load. In other words, the very mechanism that might alleviate Resource from strong capacity constraints rendered coding fidelity *less* resource-like. This conundrum points to the need for continuous report tasks with set sizes that significantly exceed estimates of capacity on categorical report tasks. Such large set sizes can have significant effects on capacity [*e.g.* 12 items (Matsuyoshi et al., 2014)] and may provide insight into its mechanistic relationship with precision. This relationship has received much less attention than each of its elements. In this regard, tremendous insight into WM capacity and precision has been gleaned from studies focusing on their differences, guided by the principles of Slot and Resource respectively. Our study suggests that their commonalities may be just as informative.

